# Functional morphological innovation corresponds to shifting lines of genetic least resistance

**DOI:** 10.1101/2022.07.30.502152

**Authors:** C. Tomomi Parins-Fukuchi, James G. Saulsbury

## Abstract

Phenotypic traits are often constrained in their evolution by the genetic and developmental map underlying them. However, the extent to which these constraints themselves evolve has not been well-characterized. Traits that are integrated are generally thought to be constrained in their ability to evolve because their shared developmental and genetic architecture causes any change in one to be mirrored in the other. Nevertheless, it is not yet clear whether correlations can constrain major episodes of phenotypic diversification over longer timescales. We reconstruct patterns in integration within nine primate lineages and model the evolution of the resulting modules in a phylogenetic context. We find that patterns in integration are generally evolvable across lineages, with apes displaying particular lability in the composition of morphological modules. This increased turnover in module composition corresponds to both divergent evolution of skeletal shape and the formation of novel complexes of locomotor behaviors and postures. While integration may play a role in constraining evolutionary innovation, its effects are dynamic, shifting as the structure of integration itself evolves during episodes of exceptional ecomorphological diversification.

## Introduction

Evaluating the trade-offs between constraint and innovation remains a fundamental goal in evolutionary biology. While perhaps the most visible thread in evolutionary research has focused on the role of adaptation in shaping the emergence of new and novel characteristics, a parallel thread has aimed to understand the role of constraints (Gould and Lewontin 1979, Gould 1980, Maynard-Smith et al. 1985). Organisms may be constrained in several ways, including by history and phylogeny, or by structural limitations imposed by physics. One source of constraint that has been of particular interest to evolutionary biologists stems from the structure underlying shared genes involved in the development of phenotypes.

Phenotypic traits are often formed through shared genes involved in underlying developmental pathways. The resulting genetic pleiotropy leads to correlations between traits that share underlying genes. For example, if two measurements from the jaw tend to covary when observed over many individuals, they are said to be “integrated” and likely possess some shared developmental and/or genetic basis (Olson and Miller 1958, Cheverud 1984, Mitteroecker 2009). While organisms frequently display integration between traits, they are generally not “fully” integrated-- i.e., not all aspects of an organism are tightly correlated with one another. Instead, sets of integrated characters tend to segregate into semi- or fully independent “modules” that, while internally correlated, remain distinct from one another in their genetic and developmental basis. One common approach through which to investigate patterns in modularity, integration, and constraint is by constructing a matrix of within-population correlations between a set of characters, known as the P-matrix (Cheverud 1988). Since trait correlations within populations stem from shared underlying genes and development, they provide an indirect view into the role of constraints in shaping phenotypic change.

Integration within modules is thought to introduce evolutionary constraint, because as one trait changes, so do all other traits with which it is correlated. As a result, if two positively correlated traits each have selective optima in opposing directions, they can only reach their respective optima if the constraint imposed by their shared genetic basis is relaxed. Constraints induced by genetic covariances may play a significant role in phenotypic evolution. Evolutionary change often appears to be constrained by the G-matrix, biasing change in the direction of least constraint (e.g., Houle et al. 2017, Sztepanacz and Houle 2019). On the other hand, several studies have also shown that, in many cases, adaptation can proceed relatively unconstrained by the orientation of the G-matrix (Beldade et al. 2002, Agrawal and Stinchcombe 2009). Nevertheless, quantitative traits do appear to show some tendency to evolve in the same direction as the genetic covariances. This pattern has been hypothesized to reflect trait means adapting along ‘lines of least genetic resistance’, although it could also result from evolution of the matrix itself (Schluter 1996). In the former scenario, adaptation is constrained in directions that align with the orientation of genetic constraints. In the latter scenario, rather than constrain the directions of adaptation, the G-matrix itself changes in response to selection, facilitating evolution of the trait means along the axis favored by selection. This possibility would amount to a remodeling of the structure of integration during times of strong divergent selection between traits. A fundamental question in this area is thus whether the primary tendency of the G-matrix is to: 1) constrain the directions taken by adaptation or 2) evolve in response to selection alongside trait means.

The ramifications of genetic and developmental integration on evolution over deeper timescales remains unclear. Most of the work examining the impact of genetic constraints on phenotypic evolution has been conducted over fairly short timescales (but see, for example, Hunt 2007, Rossoni et al., 2019). Some evidence has suggested that strong integration constrains evolutionary change, with periods of relaxed integration corresponding to higher evolvability (Young et al. 2010, Evans et al., 2017). On the other hand, some authors have argued that relaxed integration leads to *lower* evolvability due to a reduced capacity of traits to evolve at a sufficiently rapid pace (Wagner 1988, Rolian 2020). This latter perspective is further echoed by work suggesting that lower modularity and tighter integration may often facilitate rapid evolutionary change (Evans et al. 2021). A third hypothesis suggests that reorganization of the structure of integration facilitates more dramatic evolutionary change (Vermeij 1973, Wagner 2018, Parins-Fukuchi 2020). This latter view may help to reconcile apparently conflicting lines of evidence by suggesting that *changes* in the structure of integration, characterized by a temporary relaxation of module boundaries followed by reformation of new boundaries, are the decisive factor in facilitating major episodes of phenotypic change. In the case of the latter hypothesis, it would be difficult to determine whether episodes of major phenotypic change were *caused* by a temporary relaxation and restructuring of phenotypic integration, or whether such structural shifts simply *accompanied* phenotypic diversification.

In this study, we sought to address two main questions: 1) how evolvable is the structure of integration? and 2) does divergence in the structure of integration correspond to divergence in skeletal functional morphology? Addressing these two questions will shed greater light on how the evolution of integration and modularity over short timescales scales to shape the evolution of novel morphologies and ecological strategies over longer timescales. Apes provide a particularly compelling case study to understand these dynamics, because their diverse locomotor ecologies and skeletal morphologies provide a well-contained system in which to ask if substantial divergence in form and function may be facilitated by shifts in the structure of integration.

## Methods and Materials

### Data and code availability

All raw data (measurements and locomotor modes), code, supplemental methods and figures, are available on figshare (https://doi.org/10.6084/m9.figshare.20407194.v2).

### Morphological data

We acquired morphological data from Worthington (2012), who generously shared raw measurements of forelimb elements observed from individual primate specimens. We examined measurements observed from the scapula, humerus, ulna, radius, and carpal complex. Each genus contained roughly 20 replicates for each linear measurement, except for Pan, which contained roughly 40 (~20 each from *Pan paniscus* and *Pan troglodytes*). Specific landmarks and measurement methodology is thoroughly outlined in Worthington (2012). Several of the genera (e.g., Gorilla) sampled in the dataset are highly or moderately sexually dimorphic. We corrected for the resulting bimodality by subtracting the difference in male and female means from each male specimen. The raw dataset of morphological characters (both sex-corrected and uncorrected) used in the following analyses are available in the data supplement.

### Phylogeny

We used the dated phylogenetic tree of 9 haplorrhine lineages from Parins-Fukuchi (2020).

### Module reconstruction

The most common approach for examining patterns in morphological integration and modularity among sets of quantitative traits is to reconstruct a correlation matrix between the traits (the P-matrix) (e.g., Olson and Miller 1958, Rossoni et al. 2019). However, our sample sizes were inadequate (Grabowski and Porto 2017) to reliably reconstruct covariance matrices within lineages and interpret the individual correlations on a continuous scale. We therefore developed an approach to examine modularity in a discrete space. First, we reconstructed a Boolean matrix of inter-trait correlations *within each genus* by fitting linear regressions between each possible trait pair within each skeletal element (or complex, in the case of the carpus). Character pairs were set to 1 in the Boolean matrix if their correlation was statistically significant after multiple comparison correction using the Benjamini-Hochberg procedure (BH; Benjamini and Hochberg 1996) and 0 otherwise. We believed the BH correction to be most appropriate for this dataset. The more common Bonferroni correction imposes an overly conservative cutoff for significance. We were concerned that it could therefore lead to spuriously rejected correlations between traits in different genera only due to subtle differences in noise or power across genera. This could lead to spurious patterns in modular convergence due primarily to different, “patchy” patterns of false negatives across genera. Of course, rampant false *positives* between traits could lead to a similar problem. As a result, using a multiple-comparison correction, but one less strict than Bonferroni, seemed the best trade-off between these two opposing risks. We then treated this matrix as an adjacency matrix for community detection under the “Louvain” method (Blondel et al. 2008). We interpreted the resulting clusterings as the discrete pattern of modularity resulting from developmental genetics/pleiotropy for each skeletal element within each genus.

### Phylogenetic analysis of modularity

The previous analyses yielded reconstructions of the discrete structure of variational modularity within each genus and for each skeletal region. We then sought to reconstruct the evolution of skeletal modules on the phylogeny, facilitating inferences on the developmental and genetic systems that determine these modules. We developed a parsimony-based procedure to reconstruct ancestral patterns in modularity, treating the clusterings yielded in the previous step as the input data for each tip in the phylogeny. Ancestral module reconstruction involved two steps: 1) reconstructing a consensus clustering at each internal node from the clusterings at each descendant node during a post-order traversal (starting at the tips and ending at the root) and 2) setting the root to a state of complete integration and performing a new consensus at each internal node between its descendant nodes *plus* its parent node in a pre-order traversal (starting at the root and moving to the tips). This is a conservative treatment of the root, tending to bias against spurious reconstruction of trait-trait correlations at internal nodes. A full description of the algorithm can be found in the supplemental methods (also see Fig. S1).

### Measuring convergence/divergence in the structure of modularity

Using the ancestral reconstructions of skeletal modules, we identified pairs of correlated traits in each genus that were inferred to have been uncorrelated in the most recent common ancestor. We then tallied the number of convergent pairs (trait pairs that were derived independently across two lineages) between each pair of genera. To generate a reference point for comparison, we then corrected these tallies relative to the expectation under randomness. To calculate this, we designed a resampling procedure that worked as follows: assume Pan and Homo respectively had 20 and 30 derived trait pairs each among 200 possible trait pairs, 10 of which were shared in common. Our procedure would then resample vectors of 20 and 30 items from a larger pool of 200 simulated trait pairs and count how many items overlapped under this random scheme. We then repeated this procedure 100 times; the mean across all replicates was used as the expected overlap. We then subtracted the expected overlap from each raw tally of convergent pairs for each genus pair. The resulting index was zero when the number of convergent trait pairs between two genera exactly matched the random expectation, positive when the structure of integration was more convergent than expected, and negative when more divergent than expected. In addition to improving interpretability, this step also likely reduced the chances of interpreting any residual statistical errors when reconstructing trait correlations as biological patterns. Unless exceptionally strong biases in power or noise occurred in the dataset (which we have no reason to believe existed), convergence or divergence in trait integration patterns across genera should be close to zero when primarily characterized by stochastic sources of error.

### Shift in phenotypic means

We also aimed to compare shifts in modularity to changes in phenotypic means across genera. To examine broad patterns in phenotypic change as expressed by the mean phenotype for each genus, we performed a principal components analysis (PCA) on the trait means calculated for each genus. We discarded the first principal component (PC), which often primarily reflects body size information in datasets of linear measurements (McCoy et al. 2006). We then performed ancestral state reconstruction (ASR) of PCs 2 and 3 and used the resulting vectors from pointing toward each genus and originating from its most recent ancestor on the phylogeny to represent the overall magnitude (Fig. 1 and 2b) and direction (Fig. 2b).

**Figure 1.**
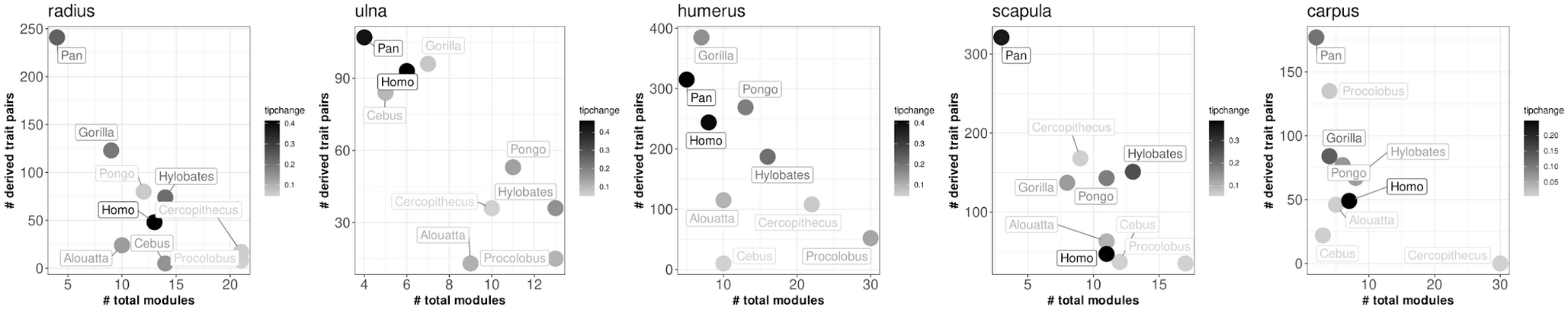
Relationship between number of modules (x-axis), the number of newly derived trait pairs (y-axis), and rate of morphological change (shading) taken from principal components analysis of the phenotypic means calculated across genera. The rate of change was calculated as the magnitude of the vectors of PC 2 and 3 change displayed in Fig. 2 divided by the length of time (in millions of years) over which the change occurred.

**Figure 2.**
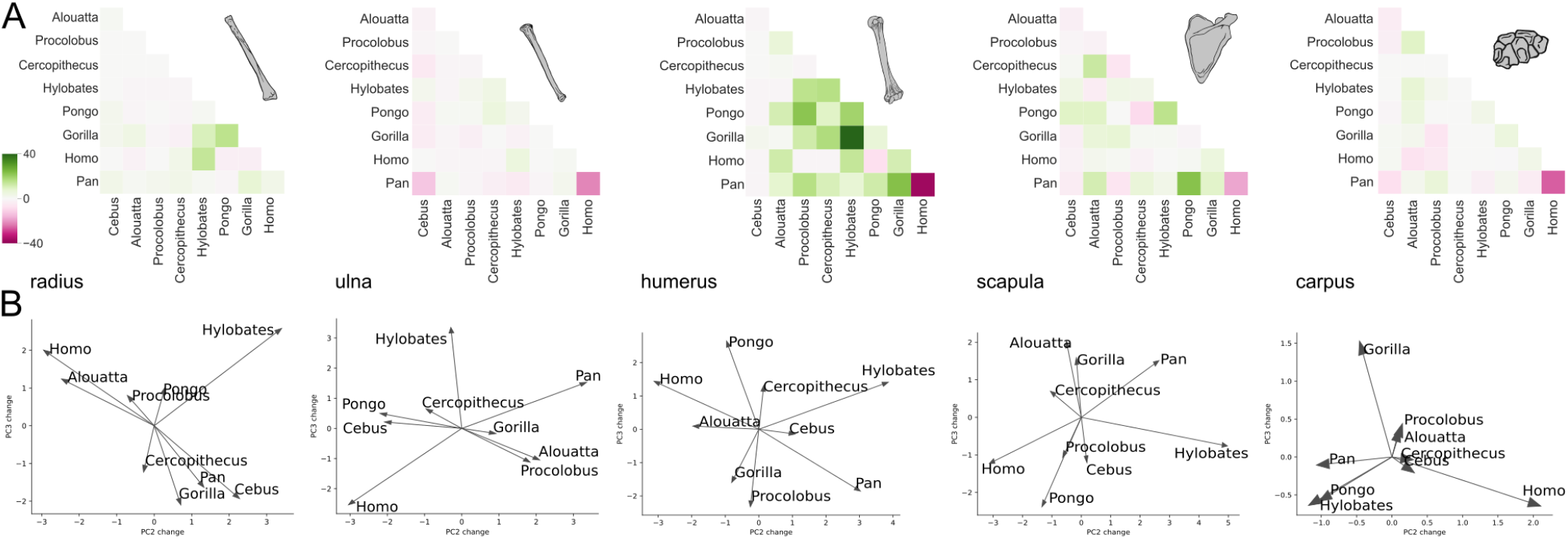
A) Number of newly derived trait pairs in the pair of genera intersecting in each cell, corrected for chance. Positive values indicate genus pairs that have each independently derived more trait pairings than expected by chance (given the number of total new trait pairs within each genus), while negative numbers indicate fewer than expected by chance. B) Vectors of genus means for PCs 2 and 3 between each genus and its most recent ancestor in the phylogeny. The anchor-point of each vector (the nearest ASR from each genus) was centered and scaled on zero such that the vectors represent the magnitude and direction of change, but not the starting and end positions in absolute PC trait space.

### Locomotor evolution

To examine coincident patterns in locomotor evolution, we also characterized the locomotor behaviors of each sampled genus. We marked as presence or absence whether each genus regularly engages in the following behaviors and postures: orthogrady, antipronogrady, brachiation, facultative bipedality, obligate bipedality, palmigrady, “fist-walking” (e.g., as observed in orangutans), knuckle walking (e.g., as observed in chimpanzees and gorillas), leaping, and bridging. Ecological behavior data and relevant references are available in the supplement. We then reconstructed ancestral states and characterized derived gains and losses of each behavior in each genus. The ancestral state reconstructions were performed using parsimony and used delayed transformation. This implies convergence in knuckle walking in chimpanzees and gorillas. Because this remains a controversial question in primate biology (e.g., Orr 2005, Begun et al. 2011), we interpreted our results in light of this controversy and the reliance of our reconstruction on the *a priori* decision of reconstruction approach.

## Results and Discussion

### Evolution of module composition

The lineages surveyed were more variable in their integration patterns than we expected, showing a fair amount of variability in module composition (see data supplement for full reconstructions made within lineages as well as ASRs). Nevertheless, most of the lineages surveyed appeared to possess a shared core of modules that were conserved across lineages, with old and new world monkeys in particular displaying high similarity due to the presence of many conserved integrated trait pairs (Fig. S2). Nevertheless, apes were more variable, with Pan in particular being highly innovative in the structure of modularity (Fig. 1). Consistent with the results of Young and Hallgrímsson (2005), we also found Hylobates to be generally less strongly integrated (i.e., display more modules) than other ape lineages. In general, while we observed more change in module composition across lineages than we expected, genera overall display a degree of conserved similarity in module composition (Fig. S1).

### Pleiotropic constraint and evolutionary innovation

Apes displayed the most consistently high rates of phenotypic change (Fig. 1). They were also more consistently innovative in the structure of skeletal modularity (i.e., possessed many derived trait pairs). Non-ape lineages occasionally derived many new trait pairs within skeletal regions (e.g., Cercopithecus in the scapula or Cebus in the ulna), but in general, apes consistently possessed many derived trait pairs across all regions. Innovation in skeletal integration patterns also appears to weakly correspond to elevated rates of phenotypic evolution. However, the correspondence is not continuous (i.e., more derived trait pairs do not linearly correspond to higher rates of phenotypic change) and the pattern is not at all visible in new and old world monkeys. The latter may be an artefact of the poor sampling of non-ape lineages. This leads to long temporal intervals between the sampled lineages, which may have led to underestimates in the rate of phenotypic change in these lineages (e.g., see Gingerich 1985). Understanding the generality of the pattern therefore requires more complete taxon sampling than was available to us. Nevertheless, the general pattern that we uncovered reveals that apes display overall high rates of change in both skeletal shape and modularity.

### Divergence in modularity and functional morphology

Pan and Homo are each highly innovative in terms of both functional morphology and the structure of integration. While apes in general are innovative in terms of locomotor behavior, both Pan and Homo display particularly unique combinations of derived behaviors (Fig. 4). In addition, despite being relatively recent sister lineages, they are strongly divergent from one another in both locomotor behavior and skeletal phenotype (Fig. 2b), with the vectors of phenotypic means directly opposing across all of the skeletal regions surveyed. The two lineages also each display many newly derived trait pairs but are significantly more divergent from one another in the structure of integration than expected by chance across nearly all regions (Fig. 2a). Thus, while each is highly innovative overall, they have undergone changes in opposing directions since their lineages diverged. This pattern demonstrates the tight link between patterns in the evolution of morphological integration and patterns in morphological and ecological divergence, while also highlighting the tendency for shifts in modularity to coincide with episodes of substantial innovation.

Changes in skeletal shape often coincide with changes in the underlying genetic links between traits, indicating that the structure of integration is itself somewhat evolvable. However, the rampant convergence observed in the integration of the humerus suggests that this lability may still be bounded. This may indicate some degree of constraint in the structure of integration. For example, while Gorilla and Hylobates are exceptionally convergent in their humeral integration patterns (Fig. 2a), they display vectors of change in skeletal shape that are uncorrelated (Fig. 2b). They are also highly divergent in terms of ecology, life history, and body size. Across all of the skeletal regions surveyed, we found a tendency for certain trait pairs to be more prone to rederivation than others. Separate regions also displayed different tendencies, with the carpus tending to re-evolve many trait pairs, and the radius encountering comparatively little convergence in modularity (Fig. 3). It is therefore possible that changes to the structure of integration are constrained such that some linkages are more readily evolvable than others, given development and the genotype-phenotype map. For example, linkages due to a non-additive phenotype map may be more readily evolvable (Rice 2004). This possibility would explain the observed pattern of rampant convergence in humeral integration that coincides with unpredictable vectors of divergence in shape evolution and differences in locomotor function (Fig. 4).

**Figure 3.**
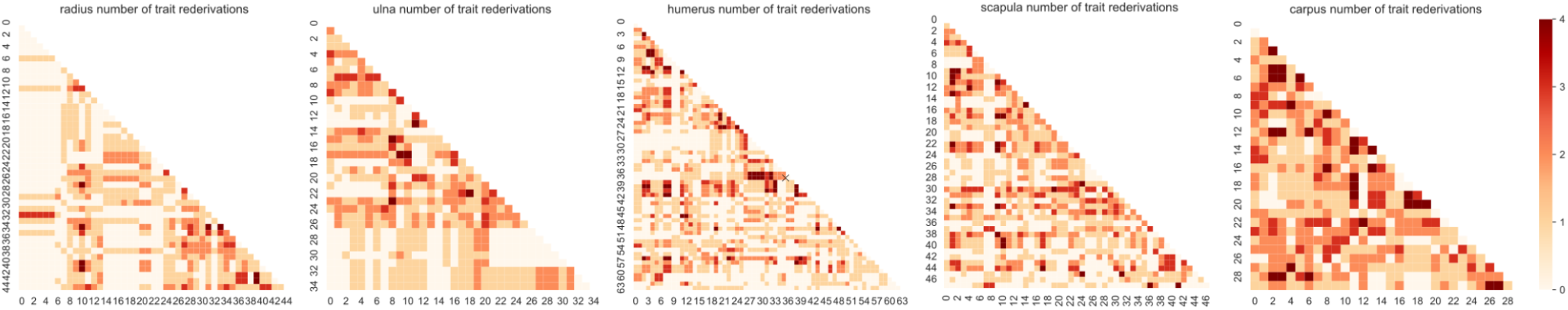
Number of times the pair of traits represented in each cell of the matrix has been uniquely derived across the tree.

**Figure 4.**
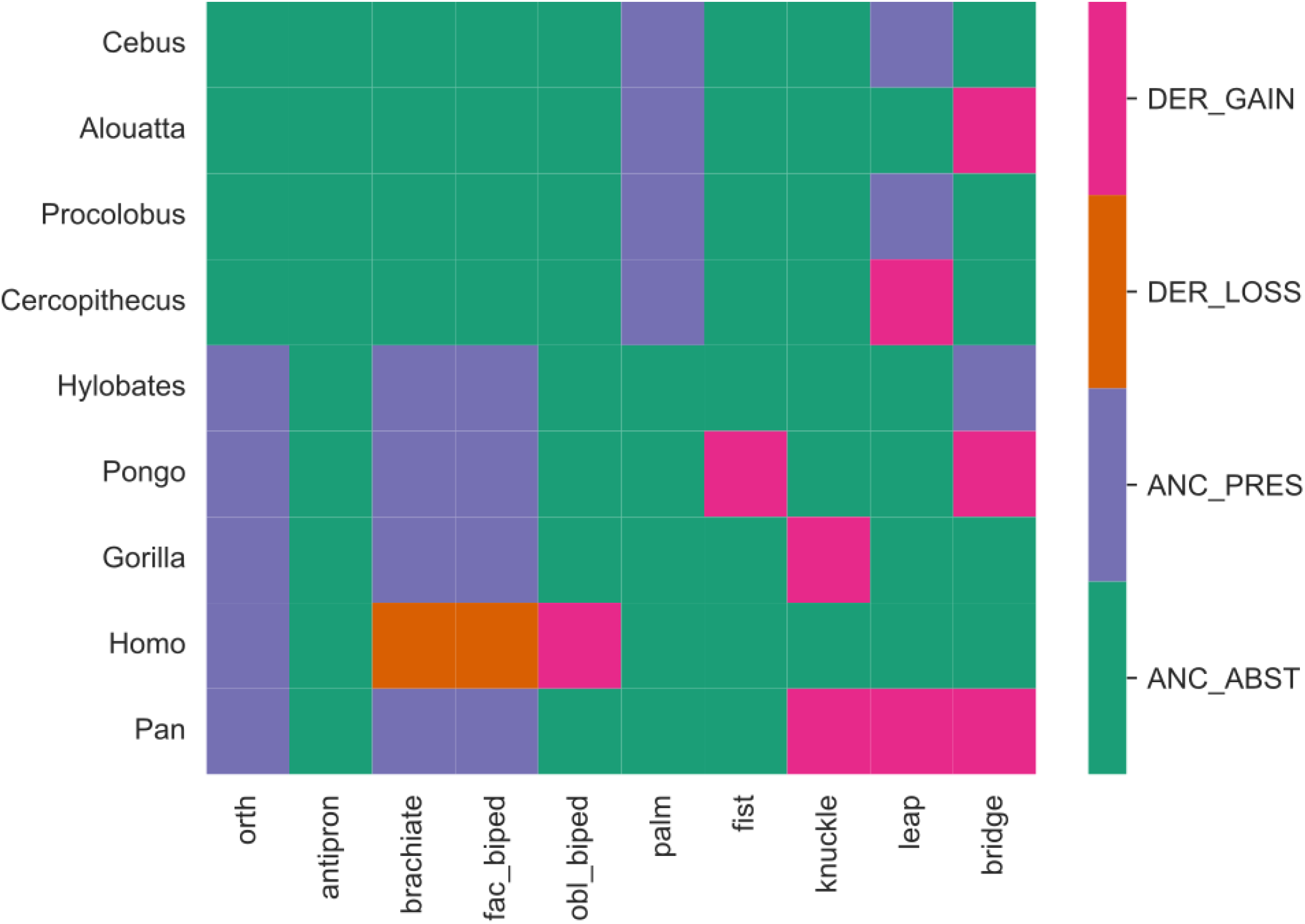
Evolution of locomotor modes across the 9 primate genera examined in this study.

### Phenotypic disparification and the evolution of genetic constraint

This work demonstrates that diversification in skeletal phenotype and locomotor function has coincided with shifts in the structure of integration throughout ape evolution. This is concordant with the “breakup-relinkage” hypothesis described by Wagner (2018) and further explored by Parins-Fukuchi (2020), wherein episodes of major morphological change coincide with rearrangements of the structure of integration. It may also be consistent with the hypothesis of Young et al. (2010) that apes experienced an escape of constraints imposed by genetic pleiotropy. While the results of Young et al. demonstrate that apes display reduced integration *between* serial homologs in the fore- and hindlimb, our results suggest that integration can remain quite high in apes *within* homologs. In other words, traits within individual homologs are often strongly integrated in great apes, but the structure of integration tends to be novel. While, for example, individual skeletal elements are often exceptionally integrated in Pan, the composition of the resulting modules is overall quite innovative. This is distinct from new and old world monkeys, which display a range of integration strengths, but tend to retain many of the same trait linkages within bones. So while they are often strongly integrated, the underlying pleiotropic structure underlying individual bones appears more free to change in apes. Overall, higher lability of the structure of integration within homologs coincides with phenotypic and locomotor innovation. One valuable avenue of future work would be to identify whether the reduced integration between serial homologs in apes also corresponds to differences in the structure of integration within each homolog. For example, does the structure of integration between traits from the humerus differ from those taken from the femur?

### Lines of least genetic resistance and the evolution of evolvability

Our analyses show that the structure of integration is relatively labile among apes, changing in concert with ecomorphological diversification. This pattern may stem from at least two possible processes: 1) Shifts in the structure of integration in apes reflect enhanced lability of developmental and genetic constraints such that ecomorphological diversification is able to proceed in novel directions due to the relaxation of constraints in different ways across lineages or 2) the structure of integration evolves as natural selection causes shifts in mean phenotype. In the first scenario, the lines of least genetic resistance shift at an increased pace, relaxing constraints in new ways and facilitating change in novel directions. In the second scenario, morphological integration does not act as a strong constraint at all, instead freely shifting in response to selection to facilitate change in directions favored by selection. We were not able to distinguish between these possible alternatives using the data available to us. Nevertheless, the relatively high lability in apes suggests that, if morphological integration (and underlying patterns in pleiotropy and development) *does* create strong constraints in general, it is one that apes have at least partially overcome. And in either case, enhanced evolvability of the structure of integration has played an important role in the diversification of functional morphologies across apes.

On the surface, our results may appear to conflict with quantitative genetic work suggesting that trait integration acts as a constraint on evolutionary change over both short and long timescales, particularly in model systems such as *Drosophila* (e.g., Houle et al. 2017, Sztepanacz and Houle 2019). Conversely, our results are consistent with other work that has found variation in the extent to which trait integration limits adaptation (Agrawal and Stinchcombe 2009). We suggest that such patterns observed in *Drosophila* may reflect interactions between both externally-imposed functional constraints and internally-imposed developmental and genetic constraints, wherein one reinforces the other over long timescales. We suggest that our results may stem from the radically different set of ecological and functional constraints imposed on apes. The set of functional morphologies displayed by and accessible to apes are quite widely variable, providing a potential impetus for the dissolution of old modules and the formation of new ones. At the same time, the recurrence of modules between genera with widely different functional morphologies also suggests some degree of constraint in the available developmental-genetic space from which new anatomical structures can be constructed.

The dynamic outlined above hints at two possible competing lines of causality underlying the relationship between changes in module composition and ecomorphological diversification. It is possible that the dramatic diversification between apes in locomotor function and skeletal morphology may have been facilitated by an increase in the lability of module composition encountered early in the divergence of apes and sustained until the present. However, we speculate that it may be more likely that causality stems from the opposite direction, with apes’ increased tendency to encounter new habitats driving the evolution of new permutations of trait modules in order to facilitate the emergence of novel skeletal morphologies and locomotor functions. At the same time, unique aspects of the ape body plan, such as orthogrady, may have additionally predisposed the increased diversification in both skeletal form and the underlying composition of modules. In any case, further study will be needed to better understand whether there is a greater tendency for releases in the structure of integration to facilitate episodes of evolutionary diversification, or whether integration tends to evolve in response to functional demands imposed by the environment and interactions between traits.

### Differing strategies to locomotor flexibility among apes

The vastly divergent skeletal phenotypes and structure of integration across the knuckle and fist walking apes (Pongo, Gorilla, Pan) support the hypothesis that these behaviors were acquired independently across all three lineages. This is because, while the behaviors are superficially similar (especially in the case of Gorilla and Pan), they appear to have distinct morphological and developmental origins. As a result, while the ancestral state reconstruction of locomotor behaviors were not equipped to support the convergence vs. shared origin of knuckle-walking hypothesis, we believe that the divergence in the structure of integration between Pan and Gorilla suggests that the behaviors may have been acquired independently, arising in distinct developmental, genetic, and perhaps ecological contexts.

Hylobates, which are both highly specialized and unique in their locomotor behavior, displayed relatively high rates of evolution in shape across most of the skeletal regions. While gibbons were not the only lineage that was less integrated overall (e.g., Procolobus and Cercopithecus across all regions besides the carpus), they did more consistently display a larger number of derived trait pairs. So while gibbons are overall less integrated than other apes, the formation of novel axes of morphological integration still appear to play an important role in their evolution. Overall, skeletal evolution in Hylobates may be best characterized as a system of comparatively relaxed integration that nevertheless retains the ability to form new modules.

### Conclusions

While our study provides only a limited examination of the evolution of the structure of morphological integration and its relationship to phenotypic and ecological diversification, it does yield several important findings. The structure of integration appears evolvable across apes, with new trait pairs being regularly derived and rederived. However this evolvability appears constrained in some dimensions, with some trait pairs breaking apart and re-forming more readily than others. Overall, the structure and overall strength of morphological integration play an important role in morphological evolution, but their effects are often unpredictable and varied. We found that apes, which have undergone a high degree of locomotor innovation, also more readily derived new trait linkages as compared to new and old world monkeys. Overall, these findings suggest that the evolution of novel functional morphologies may be facilitated by flexibility in the structure of phenotypic integration.

