## supplementary methods for "Functional morphological innovation corresponds to shifting lines of genetic least resistance"

We developed a parsimony-based algorithm to reconstruct ancestral modular patterns by reconstructing changes in discrete clusterings at each tip in a phylogeny. The procedure was performed as follows (also see Figure S1):

1) Starting from the tips (i.e., in a post-order traversal), visit each internal node. At each node, resolve a temporary ancestral pattern in modularity by creating a consensus between each of the child nodes. This consensus is performed by combining all pairs of traits that are housed within the same module in both child nodes. If a trait pair is not found in both descendants, they are placed in separate modules in the ancestor.

2) After reaching the root, set the root state to represent a state of complete integration-- i.e., all characters occupy a single shared module. An alternative procedure might assume the opposite-- that the ancestral node represents a state of complete disintegration, with each trait independent. However, this may lead to the discovery of falsely convergent modules as an artifact in cases where modular patterns are shared across all lineages. The approach taken (complete integration in the ancestor) was therefore more conservative given one of our main questions-- whether convergent trait vectors and ecological functions correspond to convergence in the structure of modularity.

3) Starting from the root, proceed toward the tips. At each node, estimate the ancestral pattern in modularity by calculating a consensus between the node's parent and each descending tip. The consensus is performed by generating an $n$ by $n$ matrix (where $n$ is the number of traits). In each cell, tally the number of times the corresponding trait pair occurs in the same module the node's parent and descendant nodes. All trait pairs that occur in 2 or more out of the 3 surrounding nodes are placed within a single module (Fig. S1b). Under some combinations of descendant module configurations, this arrangement can group traits that do not form a full clique. An alternative approach would involve arbitrarily choosing between conflicting cliques. Since our goal was to examine convergence among newly derived trait pairs, arbitrarily choosing one clique (as opposed to combining conflicting cliques) could have yielded spurious inferences of derived trait pairs. Therefore, choosing the more inclusive ancestral reconstructions was more conservative for our biological question.

*Sampling limitations*. Reconstructing the dynamics and patterns underlying the evolution of integration and modularity is a very challenging statistical problem. Our study was limited by low intra-taxon sample size, which restricted our ability to calculate detailed patterns in integration via covariance matrices. However, this remains a consistent problem shared across many studies investigating integration patterns, given the need to manually obtain morphological measurements across many lineages. Nevertheless, we feel our approach was capable of shedding light on the coarse patterns in module evolution across lineages.

**Supplemental Figures**

*Figure S1.* Algorithm for reconstructing ancestral patterns in modularity ***a)*** at each non-root internal node, combine into modules any traits that are paired in both descendants. ***b)*** after placing all traits in one module at the root,estimate ancestral modularity at each internal node as the consensus between the clusterings at the child lineages and the parental lineage.

*Figure S2.* Overall similarity of modules reconstructed between all included genera, as measured by adjusted mutual information (AMI) between each clustering at the tips. This includes similarity due to both conserved and newly derived trait pairs.
