## Supplementary figures and images for "Functional morphological innovation corresponds to shifting lines of genetic least resistance"

### figure S1

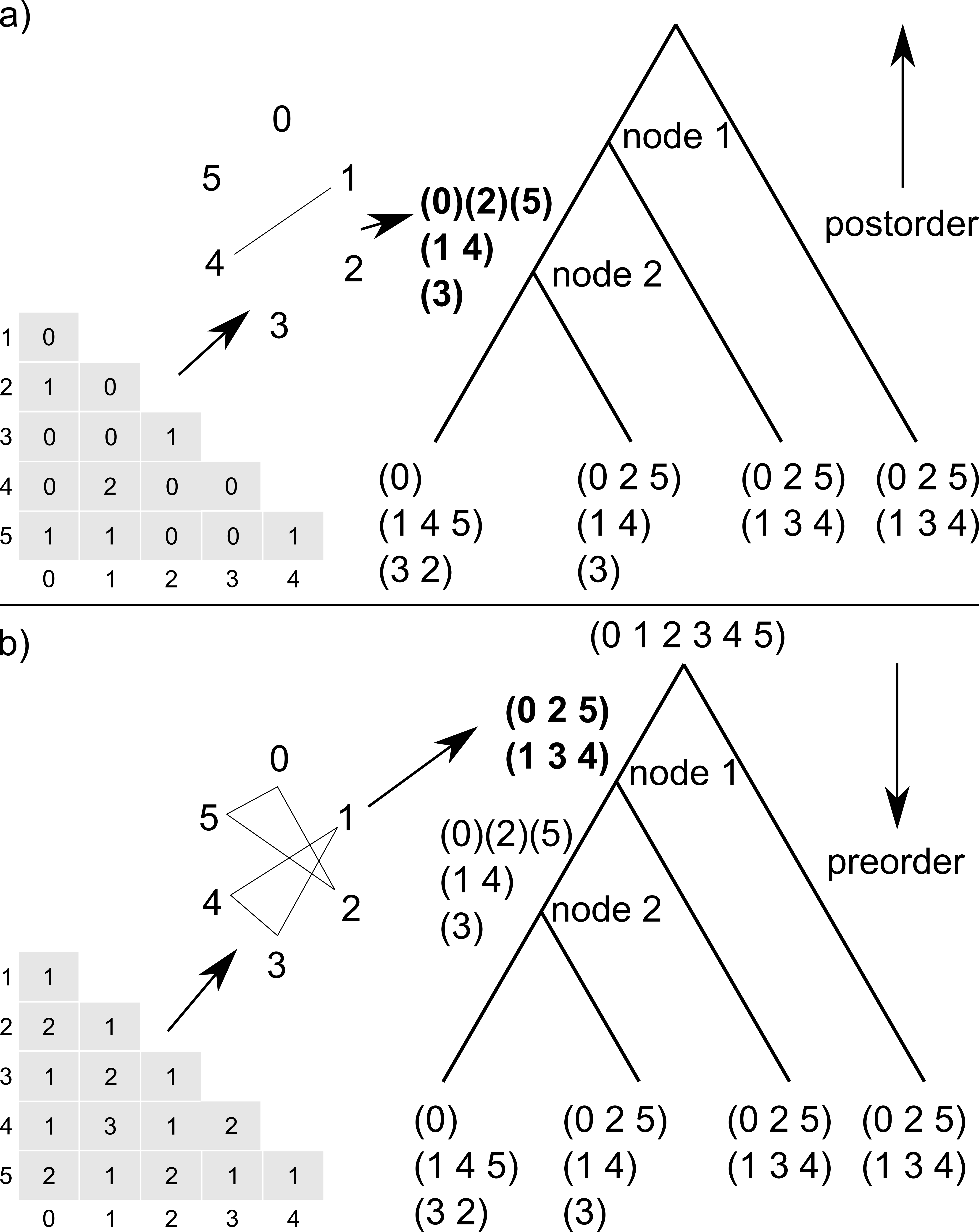

### figure S2

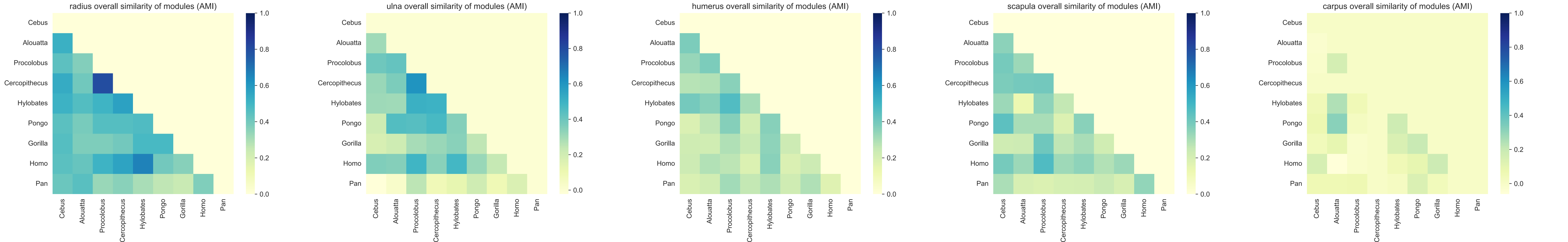
